# Scribe: next-generation library searching for DDA experiments

**DOI:** 10.1101/2023.01.01.522445

**Authors:** Brian C. Searle, Ariana E. Shannon, Damien Beau Wilburn

## Abstract

Spectrum library searching is a powerful alternative to database searching for data dependent acquisition experiments, but has been historically limited to identifying previously observed peptides in libraries. Here we present Scribe, a new library search engine designed to leverage deep learning fragmentation prediction software such as Prosit. Rather than relying on highly curated DDA libraries, this approach predicts fragmentation and retention times for every peptide in a FASTA database. Scribe embeds Percolator for FDR correction and an interference tolerant label-free quantification integrator to enable an end-to-end proteomics workflow. By leveraging expected relative fragmentation and retention time values, we find that library searching with Scribe can outperform traditional database searching tools, both in terms of sensitivity and quantitative precision. Scribe and its graphical interface are easy to use, freely accessible, and fully open source.

## INTRODUCTION

Tandem mass spectrometry (MS/MS) is the method of choice for interpreting the proteome.^1^ Since its development, data-dependent acquisition (DDA)^2^ continues as the benchmark analytical technique to detect peptides from proteolytic digests in shotgun proteomics experiments.^3^ With this method, peptides are selected and fragmented in a mass spectrometer using a variety of mechanisms that break bonds between amino acids along the peptide backbone to produce ions that aid in their identification.^4^ Different fragmentation methods generate similar ions but at different frequencies. For example, resonance collision induced dissociation^5^ (CID) and beam-type CID^6,7^ both generally produce b-type and y-type ions, but ions created by resonance CID undergo only a single fragmentation event, while ions created by beam-type CID can undergo multiple events.

Proteins in biological samples are computationally identified by matching peptide sequences in protein databases to experimentally collected mass spectra.^8^ Efficient matching is performed by either converting spectra into sequence space and performing sequence alignment (as either full length *de novo* sequences^9,10^ or sequence tags^11–13^) or by converting sequences into “unit” spectra and performing spectral correlation, typically called database searching.^14–21^ While many search engines consider the relative frequency of different ion types, few actually try to predict fragmentation patterns from peptide sequences. For example, SQID^22^ predicts fragmentation patterns from the frequency of observing pairwise amino acid cleavage patterns based on existing datasets.

Another peptide search strategy is library searching, where acquired spectra are matched to previously observed peptide spectra in libraries.^23–29^ Library searching generally detects peptides with greater sensitivity,^30^ but by focusing on previously on previously observed peptides, the smaller search space often limits the total number of peptides detected with an acceptable false discovery rate. The advent of accurate fragmentation prediction using either thermodynamics-based models^31,32^ or deep learning neural networks^33–35^ has paved the way for a hybrid approach where library searching is applied to libraries of predicted fragmentation patterns for all peptides in a protein database.^36–38^ Complexity of this workflow alongside general improvements with database searching algorithms has limited the uptake of this approach for analysis of DDA-MS data; however, searching data independent acquisition mass spectrometry (DIA-MS) datasets with predicted peptide libraries has demonstrated performance and workflow gains.^39–42^

Here we describe a new DDA library search engine called Scribe that was designed to search predicted spectra from entire protein databases. Scribe borrows scoring ideas from DIA-MS library searching designed to match peptides to noisy spectra with interference. Finally, Scribe is designed to use retention time information, when available, as is the case with most modern spectrum libraries and newer deep learning predictors such as Prosit.^34^

## METHODS

### LC-MS acquisition

Digested tryptic peptides from the Pierce™ HeLa Protein Digest Standard were separated with a Thermo Easy nLC 1200 equipped with a 15 cm Acclaim™ PepMap™ 100 C18 analytical column loaded with 2 μm beads. We used 500 ng of peptides on column for each injection. Peptides were emitted into a Thermo Exploris 480 configured to acquire DDA in a top-20 configuration with 15 second dynamic exclusion (10 ppm window, including isotopes). Precursor spectra were collected from 400 to 1600m/z at 120,000 resolution (AGC target of 250%, auto max IIT). MS/MS were collected on +2H to +5H precursors achieving a minimum AGC of 1e4. MS/MS scans were collected at 15,000 resolution (AGC target of 250%, auto max IIT) with an isolation width of 1.4m/z with a NCE of 22, 27, 32, 37, or 42. Raw files were read directly by MSFragger and MaxQuant, or were converted to mzML for Scribe and SpectraST using ProteoWizard.^43^

### Additional data sources and availability

Previously published human DDA data was downloaded from Pride PXD027242 [https://www.ebi.ac.uk/pride/archive/projects/PXD027242] (Fusion Lumos HEK 293 data) and Massive PXD017705 [https://doi.org/10.25345/C5BD2H] (QE-HF *P. falciparum* data). Exploris 480 data from the HeLa experiment presented here is available at Massive PXD037533 [https://doi.org/doi:10.25345/C5PK07624].

### Library prediction

A species-specific reviewed FASTA database for Homo sapiens was downloaded from Uniprot (2 March 2022, 20,361 entries). This database was digested *in silico* using EncyclopeDIA^44^ assuming either Trypsin, Glu-C, or Asp-N enzymes to create all possible +2, +3, and +4H peptides between 7 and 30 amino acids (matching the restrictions of Prosit)^34^ allowing for up to one missed cleavage and variable oxidation. To fit within the Prosit web server upload limitations, each charge state for each digestion enzyme was uploaded to the web server, processed individually, and combined again to form digestion enzyme-specific libraries in the DLIB format. For this work, we used either the 2020 HCD (beam-type CID) or resonance CID fragmentation model with the 2019 retention time model, where results were exported in MSP format.

While the instrument used to collect the *P. falciparum* dataset (QE-HF) was configured for NCE=27, we previously optimized the instrument to NCE=33^40^ relative to the Procal scale^45^ used to develop Prosit. Similarly, the HEK 293 instrument (Fusion Lumos) was configured to NCE=27, but optimized for Prosit to NCE=29.^46^ The Exploris 480 was configured to NCE=22, 27, 32, 37, and 42 when collecting the HeLa dataset. These values were configured to Prosit NCE=16, 29, 34, 41, and 50.

### Search engine settings

Effort was made to match settings across search engines as tightly as possible to ensure fair comparisons. Scribe was used for library searching with default settings. Briefly, searching was performed using normal target/decoy analysis with B/Y fragmentation. Mass accuracy was set for 10 PPM precursor, fragment, and library mass tolerances. Validation was performed using Percolator 3.01 filtering to a 1% peptide-level false discovery rate (FDR) threshold. When applicable, quantification on proteins passing a 1% protein-level FDR threshold was performed using retention time alignment (e.g., match-between-runs) with TIC normalization turned off.

MSFragger 3.4^21^ searches of the HeLa and HEK 293 datasets were performed inside Philosopher 4.2.1.^47^ A concatenated forward target and reverse decoy database was searched with a 10 PPM precursor and fragment mass tolerance, considering monoisotopic, +1N, and +2N precursor ions. Automatic mass calibration and parameter optimization were permitted. Protein sequences in the same Uniprot Human FASTA database were *in silico* digested assuming either Trypsin, Glu-C, or Asp-N enzymes, allowing for up to one missed cleavage. Peptides were required to have between 7-50 amino acids and range from 500 to 5000 m/z. Cysteines were assumed to be fully carbamidomethylated and methionines could be variably oxidized. PeptideProphet^48^ was used for FDR validation with the following options: “--decoy probs”, “--ppm”, “--accmass”, “--nonparam”, and “--expectscore”, which allow for additional high-mass accuracy analysis and non-parametric distribution fitting. Filtering was performed using a 1% peptide-level FDR threshold. Quantification was performed after a 1% protein-level FDR filtering, using IonQuant^49^ with default settings (10 PPM precursor mass tolerance, 0.4 minute retention time tolerance, top 3 ions) with both match-between-runs and MaxLFQ turned on.

Additional library searching of the HeLa and HEK 293 datasets was performed using SpectraST^27^ with the Trans-Proteomic Pipeline^50^ (TPP) using Petunia v6.1.0 using default settings. Briefly, mass accuracy was set for 0.01 m/z precursor tolerances. PeptideProphet^48^ was used for FDR validation using the PPM accurate mass binning option, followed by iProphet^51^ for additional FDR validation. Filtering was performed using a 1% peptide-spectrum match (PSM)-level FDR threshold but reported at the unique peptide-level, due to constraints using TPP with peptide-level FDR estimation.

Finally, MaxQuant searches of the P. falciparum and RBC datasets were performed using MaxQuant 1.6.5.0^52^ to benchmark label-free quantification. MaxQuant was configured to use default parameters with a 20 ppm fragment tolerance, where cysteines were assumed to be fully carbamidomethylated and methionines could be variably oxidized. In addition, variable protein n-terminal acetylation was allowed. Searches were filtered to a 1% peptide-level FDR and quantification was performed using unique and razor peptides with match-between-runs turned on.

### Scribe algorithmic design

Scribe runs with Java 1.8 (or higher) on Windows, Linux, and MacOS X. The general algorithmic architecture (**Figure 1**) includes a module for matching peptides to spectra based on a specified precursor mass tolerance, scoring potential peptide-spectrum and decoy-spectrum matches to produce a variety of score features, and using Percolator 3.01^53^ to combine and FDR-correct those features. If retention times are available, Scribe will create a retention time alignment from high scoring peptides and use that alignment to promote other potential peptides for a given spectrum. These newly selected matches are re-processed with Percolator and passing peptides are quantified using precursor intensity integration.

**Figure 1:**
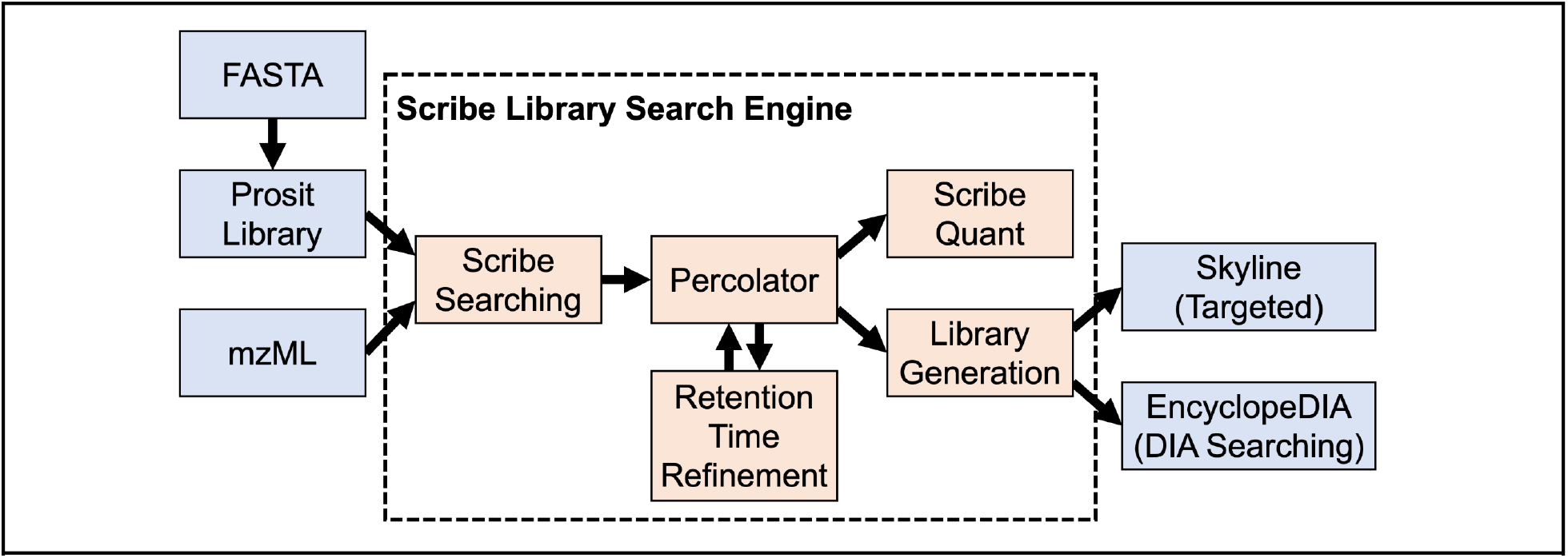
Algorithmic design for the Scribe Library Search Engine. Scribe reads mzML raw data files and spectral library files in several formats. Scribe then computes score features for peptide-spectrum matches and runs Percolator for FDR assessment. Scribe result files are quantified at the precursor level, and can be used directly as libraries in Skyline and EncyclopeDIA for future DIA or targeted experiments.

Scribe is algorithmically optimized for high resolution MS data, where both raw file data (as mzMLs) and library data are first indexed in open format SQLite databases. These indices enable rapid, multi-threaded matching between acquired and library MS/MS in randomly sized chunks, such that jobs transfer data to and from slower hard disks to faster random-access memory asynchronously. Both raw file and library indexes are saved for future searches.

### Decoy generation

An equal number of decoy library entries are generated from library entries by reversing the peptide sequence (except for the N- and C-termini) for each entry. Then the intensities of each fragment ion in the library entry that are associated with amino acids (e.g., b-type and y-type for CID as well as expected neutral losses from PTMs) are assigned a new decoy mass according to the same ion type in the decoy sequence. Ions that cannot be assigned to amino acids are preserved at the same m/z in the decoy entry.

### Key library matching scores

Before scoring is performed, the square roots of intensities from both acquired MS/MS and library spectra are calculated and used for matching. This somewhat downweights the effect of extremely intense ions and upweights the effect of matching lower abundance ions.

The primary score (Equation 1) in Scribe is a negative log transformation of the sum of squared intensity errors for matched fragment ions (*i*) between the acquired (A) and library (L) spectra, where the intensities in each spectrum are normalized to sum to 1.

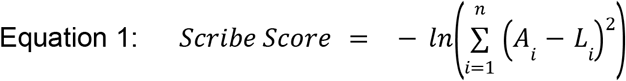

The top five matches for each acquired MS/MS are recorded and several auxiliary scores are calculated that fall into three classes: spectral similarity, precursor isotopic similarity, and mass accuracy. Several spectral similarity scores are adapted for library searching from other search engines, including XCorr and Sp from Sequest,^14^ and HyperScore and e-value estimation from X!Tandem.^16^ IsotopeDotProduct, a score from Skyline,^54,55^ calculates the correlation of precursor isotope intensities to predicted intensities based on peptide chemical structure. Other auxiliary scores are enumerated in Supplementary Table 1.

### Retention time refinement

Before refinement, scores for the top matching PSM for each acquired MS/MS are presented to Percolator for discriminant score generation and FDR estimation. The retention times for peptides that pass a 1% peptide-level FDR threshold are used for retention time alignment following the KDE non-linear alignment algorithm in EncyclopeDIA.^44^ If a peptide-spectrum match does not fit this alignment within three standard deviations (i.e., p-value<0.005) then lower scoring peptides that fit the alignment are considered for that spectrum. Peptide-level FDRs are recomputed and quantitative data is exported. Each search produces a run-specific spectrum library, but multiple searches can be globally FDR corrected and merged to create larger spectrum libraries.

### Precursor-level quantification

After searching and FDR estimation, all passing peptide matches are quantified at the precursor-level. Extracted ion chromatograms (XIC) for all precursors considering -1, 0 (monoisotopic), +1, and +2 neutrons are extracted in parallel using a configurable retention time window, typically 8x the expected chromatographic peak width. For each peptide, the elemental isotopic distribution is calculated and the acquired/expected ratio is calculated for every time point in the retention time window to check for interferences. If the acquired isotopic distribution does not match the expected distribution for expected isotopes (0, +1, +2) with intensities >50% of the maximum isotope, then excess acquired intensity in those ions is removed to maintain the correct isotopic distribution. These XICs are smoothed using the Savitzky-Golay approach^56^ and fit to a spline interpolator. To determine integration boundaries, we normalize each XIC to the apex intensity of the spline and calculate the median normalized intensity across the three expected isotopes at each retention time. This median normalized intensity approximates the peptide peak shape using all three XICs simultaneously, making it easier to assign accurate boundaries. Once the boundaries have been determined, the spline interpolated points are used to integrate the peak.

## RESULTS

Scribe is a library search engine for DDA data designed to search large spectrum libraries produced by peptide fragmentation and retention time prediction engines. Scribe is built into the EncyclopeDIA code base and released freely under the liberal Apache 2.0 open-source license. A GUI for executing Scribe and visualizing search results is available through the freely available EncyclopeDIA binary. Scribe reads raw data in mzML format ^57^ and spectrum libraries in the EncyclopeDIA DLIB format. Scribe writes search results as DLIB files, enabling rapid building of spectrum libraries for other DDA and DIA experiments. Quantitative results are also exported as tab delimited tables. Several library interconverters exist in the EncyclopeDIA software suite to convert into and out of DLIB for several different spectrum library formats, allowing Scribe-generated libraries to be used by most downstream spectrum library software.

### Evaluation with tryptic peptides

We first evaluated Scribe using a dataset of HeLa tryptic peptides analyzed using a Thermo Exploris 480. Like most modern search engines, Scribe is highly optimized for processing speed. In our tests, a full workflow of searching, retention time alignment, FDR estimation, and quantification could be typically completed in under 20 minutes per raw file on common desktop computers, or less than 20% of the time used by data acquisition. During this time, Scribe computes a Scribe Score for every peptide comparison. The Scribe Score (**Figure 2a**) is analogous to the sum of squared intensity errors between the acquired fragmentation intensities and those found in the library, where higher scoring matches indicate less error between the relative fragmentation patterns. After the top peptide candidates are determined, Scribe computes an additional 18 auxiliary scores (**Supplementary Table 1**) for the top five matches per spectrum, including relative scores like the E-Value (**Figure 2b**), spectrum matching scores like XCorr (**Figure 2c**) and precursor matching scores like the Isotope Dot Product (**Figure 2d**). The top scoring PSM features are given to Percolator to assign some peptides to spectra in a first-pass analysis. These matches help estimate expected ranges for precursor and fragment mass errors (**Figure 2e, 2f**), and relative retention time (**Figure 2g**). In particular, retention time alignment is used to select alternate peptides (within the top 5) for outlier PSMs.

**Figure 2:**
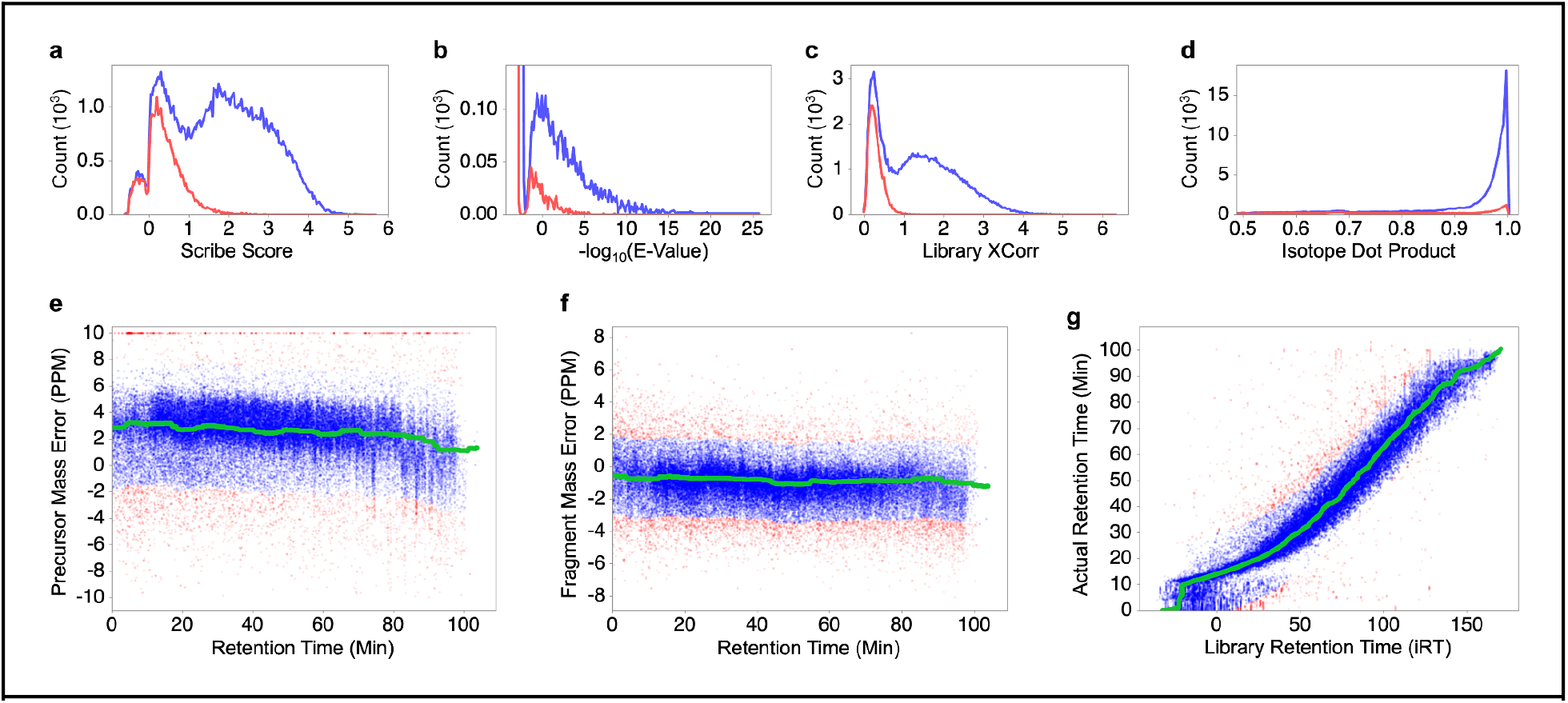
Primary scoring features in Scribe. Scribe calculates several scores for every peptide-spectrum match, including the Scribe Score (**a**), the -log_10_(E-Value) (**b**), the Library XCorr (**c**), and the Isotope Dot Product (**d**). For these scores, target histograms at the PSM-level are indicated in blue and decoy histograms are in red. In addition, global delta mass errors at the precursor (**e**) and fragment (**f**) level, as well as delta retention time (**g**) are calculated and fit using the KDE method (green line). For these error scores, the middle 90% of the target distribution are labeled as blue dots, while potential PSM outliers are labeled as red dots.

We acquired data generated from the same HeLa sample at evenly spaced normalized collision energy (NCE) settings (22, 27, 32, 37, and 42) to test the effect of library prediction quality on Scribe’s performance (**Figure 3**). Scribe (as used in this work) depends on Prosit for predicting spectrum libraries, which allows prediction tuning based on NCE. Using a panel of 1,000 peptides previously observed in HeLa, we predicted fragmentation patterns assuming integer NCEs from 10 to 50. We then compared these patterns to observed patterns in our acquired spectra and used the average of Pearson’s correlation coefficients to select the closest equivalent predicted NCE setting for each acquired NCE setting made in the method files. Interestingly, the “optimal” predicted NCE values varied from as much as -6 to +8 from the acquired NCE setting.

**Figure 3:**
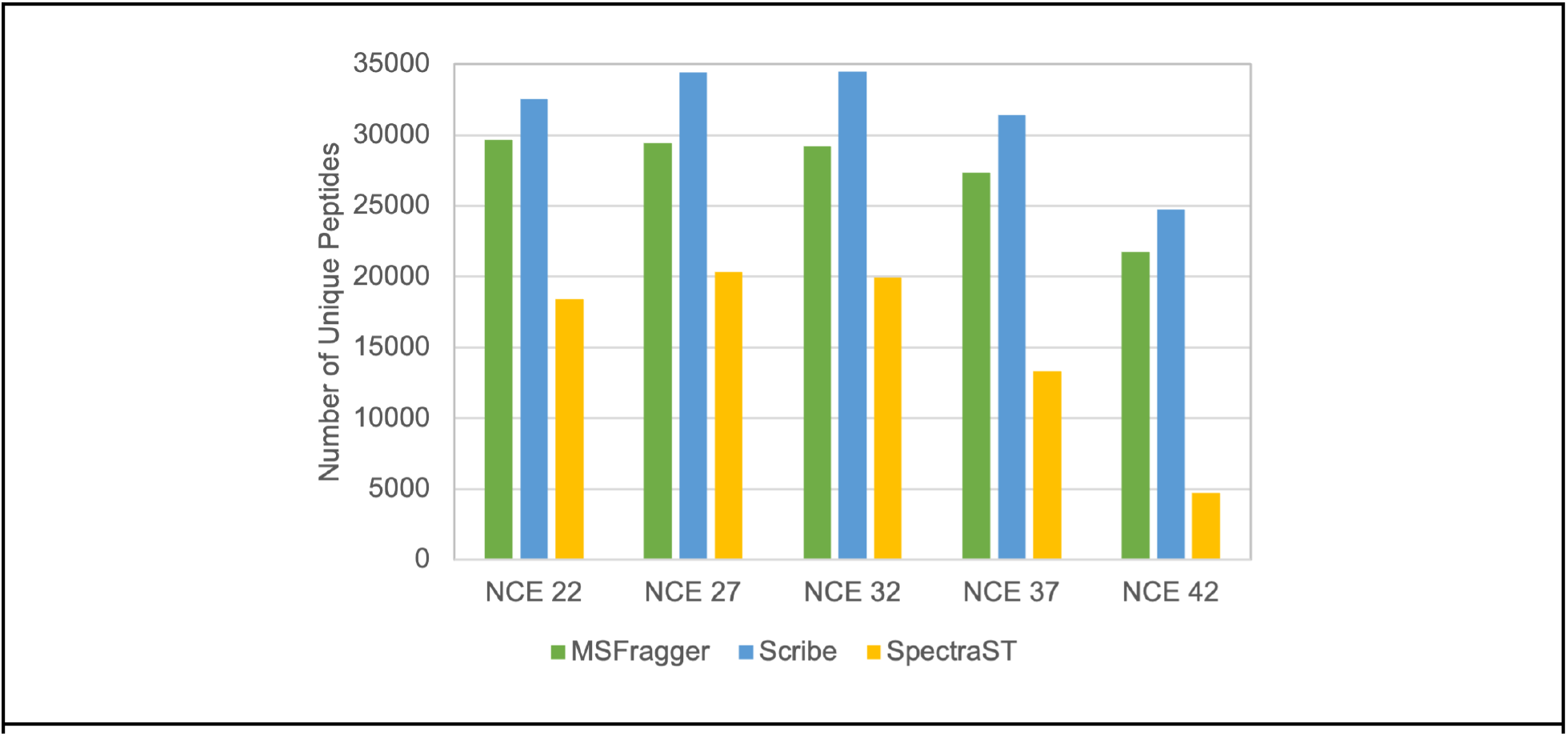
Performance with tryptic peptides. The number of unique peptides detected by MSFragger, Scribe, or SpectraST in HeLa experiments acquired on an Exporis 480 at different NCE settings. Scribe and SpectraST were provided with Prosit predicted spectrum libraries tuned specifically for the acquired NCE setting on this instrument.

While Prosit can predict peptides at NCE settings from 10 to 50, it was only trained on data collected from 23 to 38 (using the Procal scale). This means that the match quality should decrease at extreme NCEs that essentially require more extrapolation, which we observed. That said, across NCE settings in this experiment, Scribe coupled with Prosit produced 10% to 18% more peptide detections than MSFragger 3.4 within a 1% peptide-level FDR threshold. Both methods outperformed SpectraST, a current state-of-the-art library searching tool, particularly at extreme NCE settings.

Building predicted libraries with the correct NCE setting for each dataset is important. Moreover, different instruments appear to produce different fragmentation profiles at the same NCE setting, requiring NCE tuning relative to Prosit. We found that in general, searching data with a closely tuned predicted library produced the highest number of unique peptides (**Figure 4**); we found that searching with a library configured for a different NCE setting frequently performed near to, or worse than MSFragger. However, we found that in some cases, searching with an “un-tuned” library produced somewhat better results than the appropriately tuned library, particularly at extreme NCEs. This indicates that additional optimization of NCE with Prosit specifically for scoring (rather than spectrum correlation) may improve overall detection results with Scribe.

**Figure 4:**
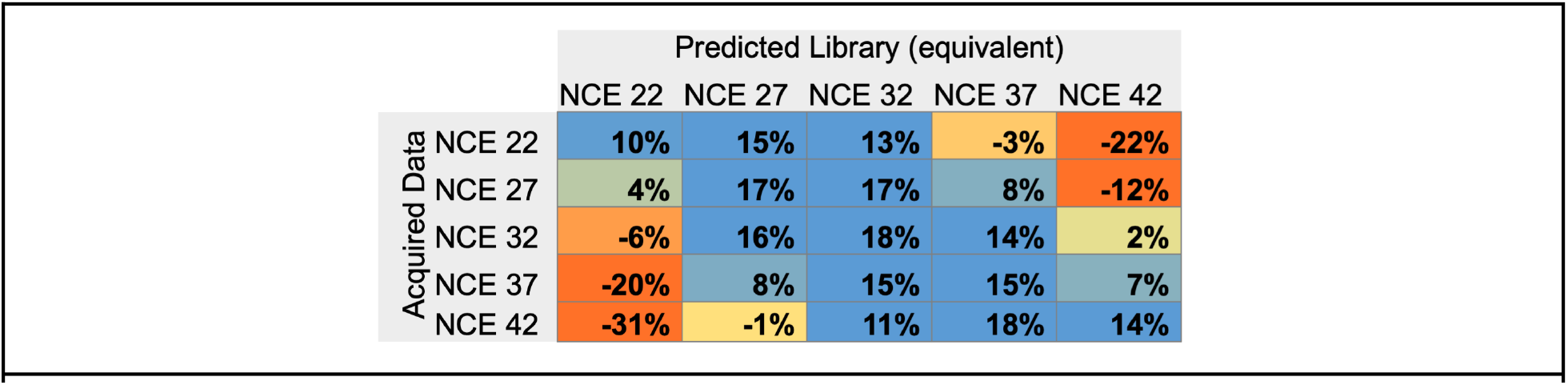
The effect of NCE on Scribe performance. The increased (blue) or decreased (red) percentage in Scribe-detected peptides compared to MSFragger depends on the NCE setting used to predict the spectrum library. Predicted NCE values are normalized to the Exploris 480 instrument used in this work. In general, tuning NCE to the instrument used to acquire the dataset results in increased performance.

### Evaluation with non-tryptic peptides

Database searching typically performs better with tryptic peptides than with non-tryptic peptides. As such, we investigated Scribe’s ability to analyze non-tryptic peptides with either beam-type or resonance CID fragmentation using DDA datasets from Richards et al 2022.^58^ In this dataset we observed similar performance for tryptic peptides, but non-tryptic peptides performance depended strongly on the digestion enzyme used (**Figure 5**). In this dataset, beam-type and resonance CID performance followed similar trends, no matter what enzyme was used.

**Figure 5:**
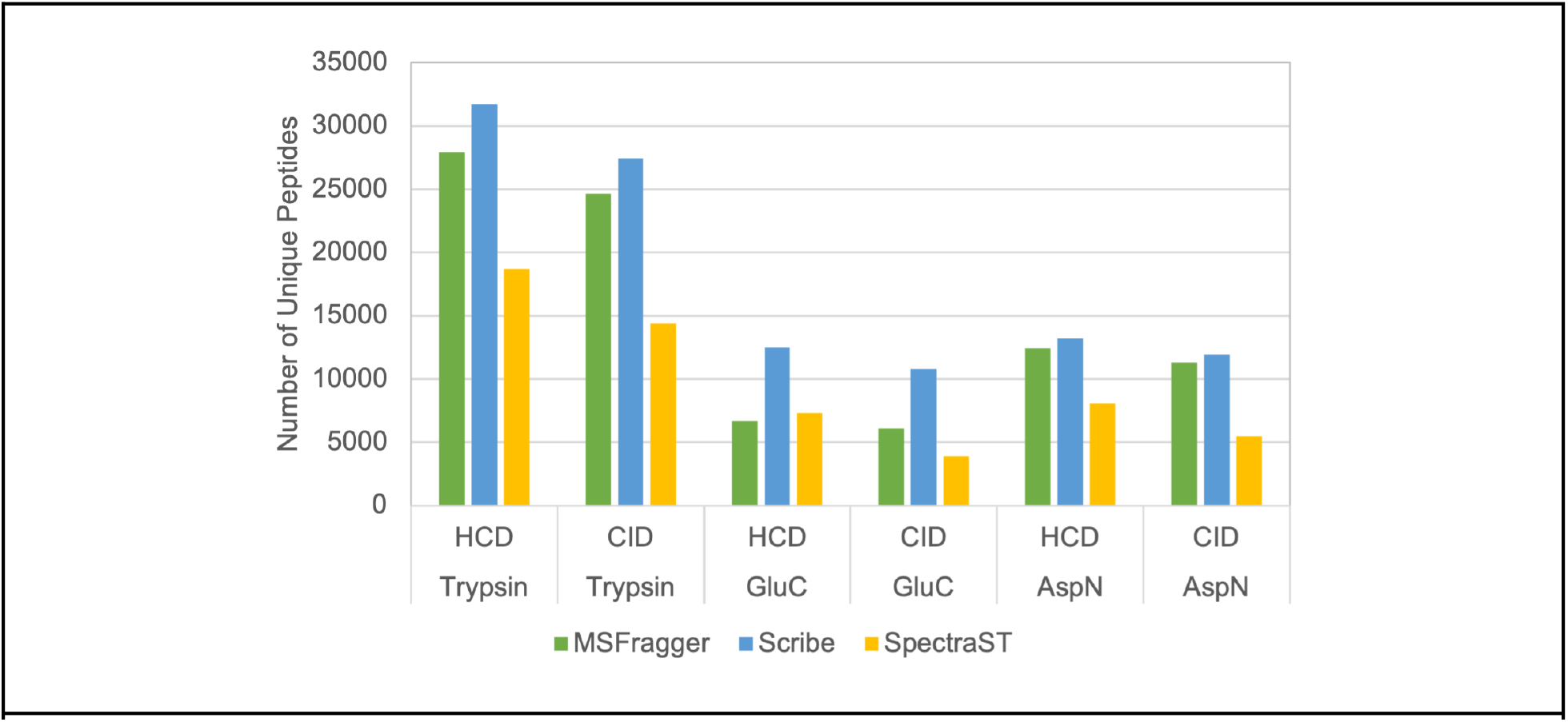
Performance with peptides derived with alternative digestion enzymes. The number of unique peptides detected by MSFragger, Scribe, or SpectraST in HEK 293 experiments^58^ with different digestion enzymes acquired on a Fusion Lumos with either HCD (beam-type CID) or CID (resonance CID).

### Quantification with Scribe

Since Scribe uses precursor-level features as part of its scoring system, it automatically calculates precursor peak areas for quantification. While integrating extracted precursor ion chromatograms is a common label-free proteomics approach, Scribe employs new ideas for interference rejection. First, Scribe calculates the expected isotopic distribution and matches it to the acquired distribution of every precursor-level spectrum in a limited retention time window. Scribe was designed to assume that it is possible to add to the expected signal through interference, but under most circumstances, it is impossible to decrease the expected signal. To account for this, if a common isotope (calculated to be >50% of the highest isotope) is observed with low intensity relative to the other isotopes, the other isotopes are assumed to have contribution from an interfering peptide. In this case, signal from the interfered isotopes is subtracted until the acquired isotopic distribution matches the calculated distribution. Coupled with match-between-runs using non-linear retention time fitting, this approach should ensure quantitative accuracy in low abundance peptides, which are more likely to experience interference.

We tested Scribe’s quantification accuracy using DDA data from a previously published matched-matrix^59^ controlled experiment of *P. falciparum* infected red blood cells (RBCs) diluted into uninfected RBCs at known ratios.^40^ Using a Prosit predicted library combining peptides from both Human and *P. falciparum* FASTA databases, we filtered search results to a 1% protein-level FDR and found that we could quantify parasite-derived peptides in up to 1:99 dilution with RBCs (**Figure 6**). In this lowest dilution, Scribe was only able to detect 338 parasite-derived proteins (plus an extra 396 RBC-derived proteins), but with retention time alignment, was able to quantify a full matrix of 1,792 total proteins with non-zero values. In contrast, MSFragger quantified a full matrix of only 681 parasite-derived proteins with non-zero values across all samples. Similarly, MaxQuant only quantified 1,295 parasite-derived proteins in the top dilution sample and failed to quantify any parasite-derived peptides in the 1:99 dilution sample, even with match-between-runs enabled.

**Figure 6:**
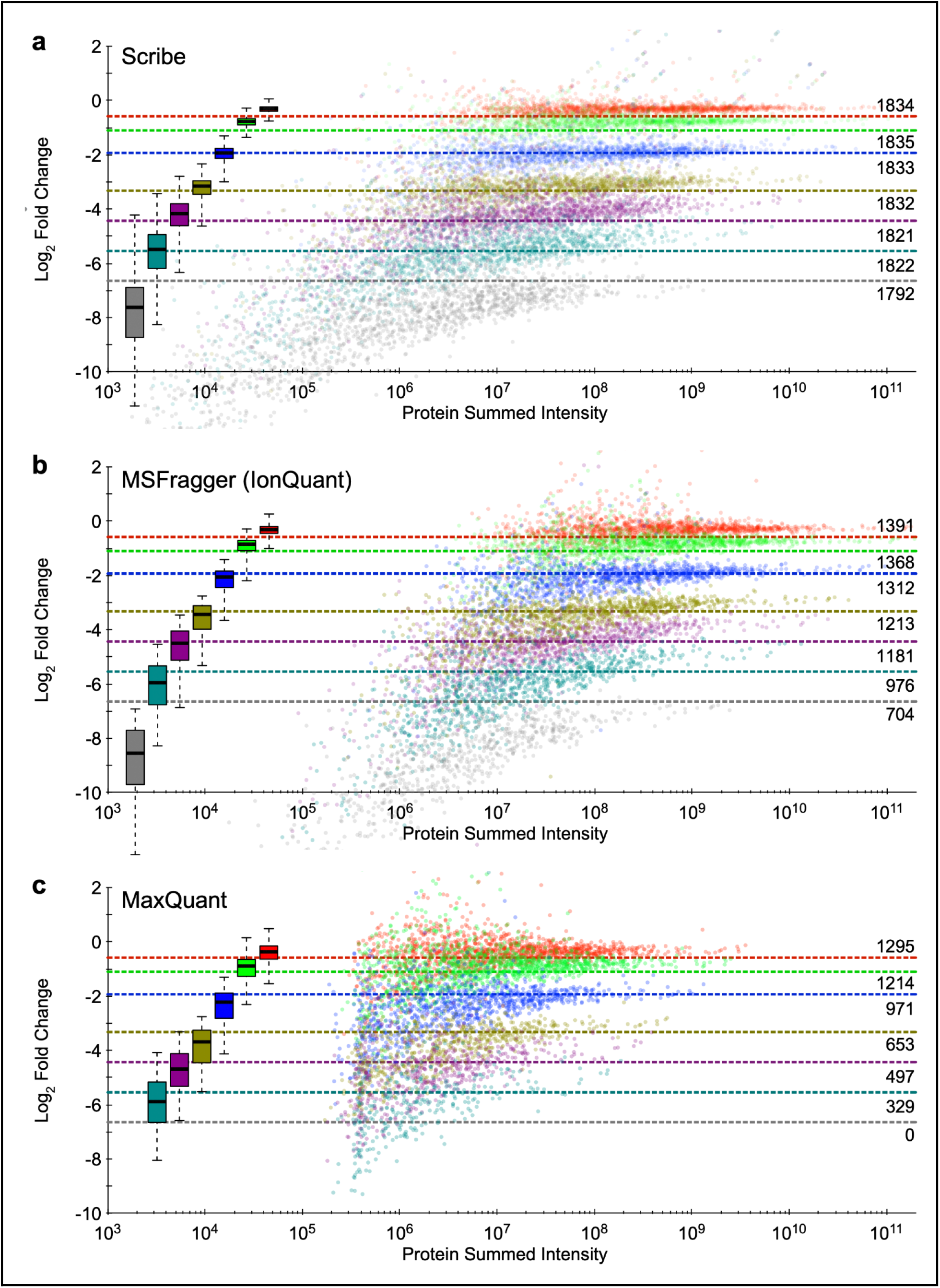
Quantification of *P. falciparum* proteins in a relative dilution with red blood cell lysates using either (**a**) Scribe, (**b**) MSFragger using IonQuant, or (**c**) MaxQuant. Seven different dilution ratios (red=2:1, green=7:8, blue=4:15, gold=1:9, purple=2:41, cyan=2:91, and gray=1:99) are shown relative to the protein intensity (summed from peptide intensities). Dashed lines indicate the expected ratio, while the number of non-zero/non-missing protein-level measurements in each dilution ratio are indicated at the right. No *P. falciparum* proteins were detected or quantified in the 1:99 dilution sample using MaxQuant. Box plots show the spread of measured values where the whiskers indicate 5% and 95% points and the bold line indicates the median measurement.

Precision of the expected ratios was also improved throughout the dilution curve, as shown by smaller interquartile ranges and 5% to 95% whiskers at equivalent expected ratios as compared to MSFragger and MaxQuant. Interestingly, due to this increased precision, we observed minor deviations from the expected ratios in the most abundant dilution samples. Since these deviations were in the same direction as those observed with MSFragger and MaxQuant, we believe they are likely caused by minor pipetting errors. Aside from these two samples, Scribe demonstrated increased accuracy in the median quantitative ratios throughout the dilution curve, which suggests that interference rejection can help improve quantitative accuracy as well as precision.

## CONCLUSIONS

When paired with predicted spectrum libraries, library searching is highly competitive with database searching in terms of sensitivity, quantitative accuracy, and speed of processing. In this work, we have strived to make Scribe easy to use, freely accessible, and fully open source.

Scribe can be used as a stand-alone search engine with automatic FDR assessment and label-free quantification. However, we feel that Scribe can also improve proteomics data analysis workflows in other areas of research. In particular, we believe that Scribe has utility as a DDA library generation tool for follow up DIA experiments. Scribe search results are immediately compatible with EncyclopeDIA as native libraries, and can be easily converted for follow up use in Skyline,^55^ OpenSwath,^60^ Spectronaut,^61^ and other tools. Additionally, the use of library spectra for detection may aid in limited material proteomics experiments, such as with single-cell proteomics.

## SOFTWARE AVAILABILITY

Scribe and an associated manual/walkthrough for how to operate Scribe is available as part of the EncyclopeDIA project: https://bitbucket.org/searleb/encyclopedia/

The version of Scribe used in this work can be directly downloaded here: https://bitbucket.org/searleb/encyclopedia/downloads/scribe-2.11.14-executable.jar

## ACKNOWLEDGEMENTS

This work is supported in part by National Institutes of Health Grant R01-GM133981 and U19-AG065156. D.B.W. is additionally supported by K99-HD090201.

---

**Supplementary Table 1:**

| # | Score Name | Score Description |
| --- | --- | --- |
| 1 | averageParentDeltaMass | Average mass error of matched precursors (including isotopes) in PPM |
| 2 | averageFragmentDeltaMasses | Average mass error of matched fragments in PPM |
| 3 | DotProduct | $\sum_{i=1}^n (A_i * L_i)$ |
| 4 | contrastAngle | $1 - (2 * \text{acos}(\text{DotProduct}))/\pi$ |
| 5 | logit | $\log(\text{DotProduct})/(1 - \text{DotProduct})$ |
| 6 | primary | $\log(\text{DotProduct}) * \text{!(numberOfMatchingPeaks)}$ |
| 7 | xCorrLib | XCorr comparing against Sequest normalized library spectrum |
| 8 | xCorrModel | XCorr comparing against Sequest model spectrum |
| 9 | scribeScore | $-\ln\left(\sum_{i=1}^n (A_i - L_i)^2\right)$ |
| 10 | numberOfMatchingPeaks | Total number of matching peaks between library and acquired spectra |
| 11 | isotopeDotProduct | Normalized DotProduct of acquired and predicted precursor isotope intensities |
| 12 | percentBlankOverMono | Precursor isotope-1 intensity divided by the monoisotopic intensity |
| 13 | numberPrecursorMatch | Number of precursor isotopes with >0 intensity (up to 3) |
| 14 | logSp | Log<sub>10</sub> of the Sp score from Sequest |
| 15 | maxLadderLength | Maximum connected fragment ions in a ladder |
| 16 | eValue | E-value of scribeScore as calculated by the Fenyo et al 2003 method |
| 17 | deltaScore | $(\text{bestScribeScore} - \text{secondBestScribeScore}) / (\text{bestScribeScore} + 10)$ |
| 18 | numConsidered | Total number of candidate library entries considered |
| 19 | chargeMatch | 1 if acquired charge matches library entry, 0 if mismatch |

